# Similar sensorimotor transformations control balance during standing and walking

**DOI:** 10.1101/2020.09.30.320127

**Authors:** Maarten Afschrift, Friedl De Groote, Ilse Jonkers

## Abstract

Standing and walking balance control in humans relies on the transformation of sensory information to motor commands that drive muscles. Here, we evaluated whether sensorimotor transformations underlying walking balance control can be described by task-level center of mass kinematics feedback similar to standing balance control. We found that delayed feedback of center of mass position and velocity, but not local feedback of joint positions and velocities, can explain reactive ankle muscle activity and joint moments in response to perturbations of walking across protocols (discrete and continuous platform translations and discrete pelvis pushes). Feedback gains were modulated during the gait cycle and decreased with walking speed. Our results thus suggest that similar task-level variables, i.e. center of mass position and velocity, are controlled across standing and walking but that feedback gains are modulated during gait to accommodate changes in body configuration during the gait cycle and in stability with walking speed. These findings have important implications for modelling the neuromechanics of human balance control and for biomimetic control of wearable robotic devices. The feedback mechanisms we identified can be used to extend the current neuromechanical models that lack balance control mechanisms for the ankle joint. When using these models in the control of wearable robotic devices, we believe that this will facilitate shared control of balance between the user and the robotic device.

## 1 Introduction

Standing and walking without falling seem to be simple motor behaviors even in uncertain environments. This is remarkable, given the instability of the human skeletal system. Continuous adaptations of muscle activity are needed to control the relatively high position of the center of mass (COM) above a small base of support. This is achieved through sensorimotor transformations: the nervous system continuously receives sensory inputs, which are processed to generate descending motor commands to muscles [1]. These sensorimotor transformations cannot be explained by local reflexes alone but rely on supra-spinal processes that integrate sensory information from multiple sources to derive information relevant to the motor task, i.e. stabilizing the musculoskeletal system [2, 3]. Experiments involving external mechanical perturbations of standing have found evidence of task-level feedback from COM kinematics. Delayed feedback from COM kinematics can explain reactive muscle activity and changes in joint moments in response to perturbations during standing [4–6]. Here we will investigate if, similar as in standing balance, COM position and velocity feedback can describe sensorimotor transformations during walking.

In contrast to standing, sensorimotor transformations controlling balance during walking are less well studied. Prior studies mainly used experimental techniques to describe the postural strategies during perturbed walking and found that subjects modulate muscle activity [7–10] to adjust foot placement (stepping strategy) [11] and the location of the center of pressure (COP) in the stance foot [8, 12]. Adjustments of COP location in the stance foot are mainly achieved through modulating the ankle moment and are therefore referred to as ankle strategy [8, 12, 13].

Following frontal plane perturbations, balance is mainly controlled using a stepping strategy. Medio-lateral foot placement is strongly correlated with COM position and velocity [11, 13–16]. Furthermore, reactive bi-lateral gluteus medius activity, which has an important contribution to foot placement, is also correlated with COM kinematics [14, 17]. These results suggest that COM kinematics are indeed important task-level variables driving the control of medio-lateral foot placement during perturbed walking [18].

Following sagittal plane perturbations, both stepping and ankle strategies are used to control balance with the latter being the dominant strategy. COM kinematics feedback can explain fore-after foot placement [15]. However, the correlation between COM kinematics and foot placement is weaker in the sagittal than in the frontal plane [19], which can be attributed to the higher reliance on the ankle strategy. Indeed, when eliminating the ankle strategy through stilt walking, a strong correlation between COM kinematics and foot placement is observed in the sagittal plane [20].

Is is currently unclear if, similar as in perturbed standing, delayed feedback of COM kinematics feedback also underlies the ankle strategy in perturbed walking. This can be evaluated at the level of both reactive ankle muscle activity and ankle joint moments. While the reactive muscle activity reflects only the neural control, the reactive ankle moment is the result of both neural control and intrinsic mechanical contributions arising from visco-elastic properties of passive tissues and active muscle fibers. Consistent with the finding that control of task-level variables rather than joint-level feedback underlie reactive standing balance [3], we expect that reactive ankle muscle activity and ankle joint moments during walking can be described by delayed linear feedback of COM position and velocity but not by delayed feedback from local joint angles and velocities.

In contrast to standing, body configuration and intrinsic mechanical stability vary considerably during the gait cycle and with gait speed. Therefore, we expect that, in contrast to standing balance, task-level feedback from COM kinematics driving muscle activity is not constant but varies throughout the gait cycle. This agrees with previous observations that the effect of a motor command (i.e. muscle excitation) on body movement strongly depends on the phase in the gait cycle [10, 21]. Therefore, we hypothesize that sensorimotor gains, describing the linear relation between COM kinematics and muscle activity, are modulated during the gait cycle and with gait speed. Experimental observations support this hypothesis. First, it has been shown that local reflexes, assessed through H-reflexes, are modulated during the gait cycle [22] and with walking speed [23]. Second, reactive muscle activity changes when applying mechanical or sensory perturbations at different instances in the gait cycle [1, 7, 9, 10]. For example, large changes in reactive calf muscle activity are observed during mid-stance [9, 10] but not during early stance or swing [10]. It has however not been investigated whether COM kinematics feedback with phase-dependent gains can explain these changes in reactive muscle activity.

If COM kinematics feedback is indeed modulated during the gait cycle, this raises the question which sensory mechanisms underly this modulation. Cutaneous stimulation studies suggest that sensory information from tactile sensors in the foot modulates reflex gains during walking [24, 25]. We therefore hypothesize that tactile sensors also modulate task-level feedback gains and thereby contribute to the observed modulation of reactive muscle activity.

In this study, we investigate the sensorimotor transformations underlying the ankle strategy to control balance during walking. We hypothesize that, comparable to standing, feedback from COM kinematics, and not local joint-level feedback, can explain reactive ankle muscle activity and ankle joint moments in perturbed walking. However, given the changes in the stability of the musculoskeletal system and changes in body kinematics during the gait cycle and with walking speed, we hypothesize that feedback gains will be modulated during the gait cycle and with gait speed based on input from tactile sensors. To test these hypotheses, we evaluated the relation between the inputs of the feedback (delayed COM and ankle joint kinematics) and the resulting motor command (muscle activity and joint torques) in four available datasets of perturbed standing and walking at different walking speeds in young healthy adults. We used different perturbation modalities, support surface translations and pelvis pushes, to excite the (neuromechanical) system in multiple ways (Fig. 1), thereby guaranteeing the generalizability of our findings.

**Figure 1:**
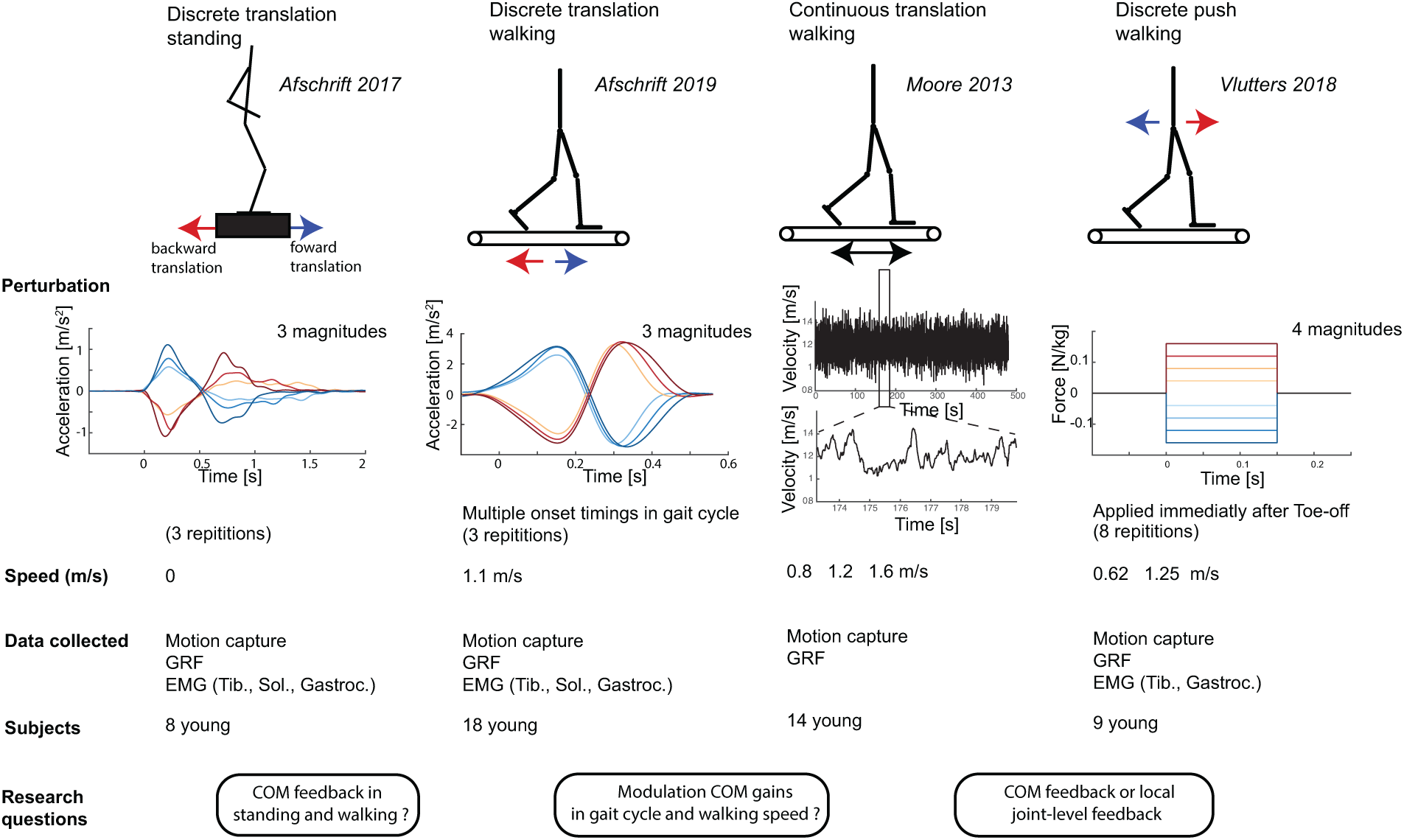
*Overview datasets and hypotheses:* We answered three research questions related to sensorimotor transformations underlying balance control by combining four previously published datasets with motion capture data of unperturbed and perturbed standing and walking. We used data of surface perturbation in standing [26] and walking [9, 10, 27] to evaluate if the ankle strategy is related to deviations in COM kinematics. Balance was perturbed during walking using both discrete and continuous surface perturbations [10, 27] and discrete pelvis push perturbations [9]. The potential modulation of feedback gains during the gait cycle was evaluated using a dataset with continuous surface perturbations during walking [27] and a dataset with perturbations applied at discrete instances of the gait cycle [10]. Finally, we used data with discrete surface translations in standing and walking to evaluate if altered sensorimotor transformations can explain the adjusted kinematic strategies to control balance observed in older adults. In all these experiments combined, joint kinematics and ground reaction forces were measured in 41 young subjects in total.

## 2 Results

In short, we showed that (1) sensorimotor transformations underlying the ankle response to perturbations of standing and walking can largely be explained by task-level feedback of COM kinematics and not with local feedback of joint kinematics and (2) COM kinematics feedback gains are modulated within the gait cycle and with walking speed.

### 2.1 Model for task-level feedback of COM kinematics

We evaluated the relation between COM kinematics and ankle muscle activity and joint moment in perturbed standing and walking. Therefore, we tested whether reactive muscle activity and reactive joint moments could be explained by a linear combination of the delayed deviation of COM position and velocity from the unperturbed reference trajectories.

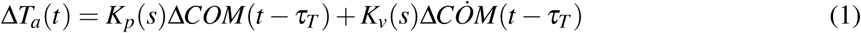

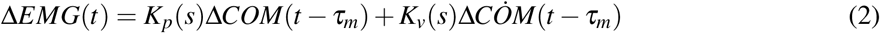

with Δ*T*_*a*_ the reactive ankle joint moment, Δ*EMG*(*t*) the reactive ankle muscle activity, *K*_*p*_(*s*) the position gain, *K*_*v*_(*s*) the velocity gain, and Δ*COM* and 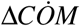 the deviation of the the whole body COM position and velocity. Note that the feedback gains (*K*_*p*_(*s*) and *K*_*v*_(*s*)) depend on the phase in the gait cycle (*s*). A neural delay of 60ms was used for muscle activity (*τ*_*m*_) and 100ms for joint moments (*τ*_*T*_). The delay in muscle activity is caused by the time needed for signal transmission and sensory integration in the nervous system [3], whereas the delay in joint moments is larger due to the additional electromechanical delay between muscle excitation and the development of force in the muscle. The sensitivity of the results to the time delay is discussed in the appendix (S1 Fig. 11).

We compared the ability of the COM feedback model to explain experimental data with an alternative model based on delayed feedback of ankle angle and angular velocity.

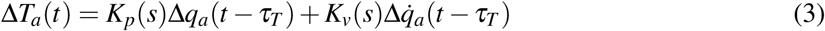

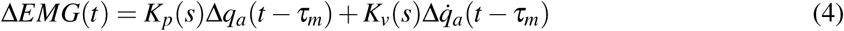

with Δ*q*_*a*_ and 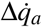 the deviation of the ankle joint angle and velocity from the unperturbed reference trajectory. A shorter neural delay of 40ms was used for muscle activity (*τ*_*m*_) and 80ms for joint moments to model the shorter latency of local reflex as compared to centrally mediated feedback pathways.

Feedback gains were estimated from the measured kinematics, joint moments and muscle activity by solving a least squares regression. Inputs (COM kinematics or ankle kinematics) and outputs (ankle moment) were selected respectively at 150ms and 230-250ms after perturbation onset since large deviations in COM kinematics were observed at this time instance for all different types of perturbations (see methods section for details). Similarly, feedback gains for the ankle muscles were estimated based on COM kinematics and muscle activity respectively at 150ms and 210ms after perturbation onset. Feedback gains were estimated for each subject individually and with the data pooled over all subjects to evaluate if subjects use similar feedback gains.

Uncentered coefficients of determination (*R*^2^) and Root Mean Square Errors (RMSE) of the measured and reconstructed joint moments, as well as muscle activity are reported to quantify the fit of the linear models. Similar as in [28], all quantities and results are non-dimensionalized using COM height during quiet standing (*l*_*max*_), the gravitational acceleration (*g*) and body mass (*m*). COM positions were normalized by *l*_*max*_, speeds by 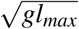, torques by *mgl*_*max*_ and muscle activity by maximal voluntary contraction values in the standing data and by maximal activity observed during unperturbed walking in the walking data (maximal voluntary contraction was not available in the walking data).

### 2.2 COM feedback explains ankle muscle activity and moment in perturbed standing

In agreement with previously published observation [5, 29], we found that reactive ankle joint moments and muscle activity can be explained by delayed feedback of COM kinematics. Delayed feedback of COM position and velocity can explain the change in ankle joint moment (*R*^2^ = 0.94, *RMSE* 0.008) and muscle activity (*R*^2^ between 0.62 and 0.89,) in response to forward and backward perturbations of different amplitudes (Fig. 2 and table 1). Tibialis anterior activity increased proportional to COM position and velocity in response to a backward directed perturbation and gastrocnemius activity increased proportional to COM position and velocity in response to forward directed perturbations (Fig. 2). Feedback gains for the ankle moment, *R*^2^ and RMSE values were similar when computed for individual subjects or pooled over all subjects, indicating that different subjects use similar feedback gains (table 1). Ankle moments reconstructed using COM kinematics were more similar to the measured data than ankle moments reconstructed using ankle joint kinematics (table 1).

**Table 1:**
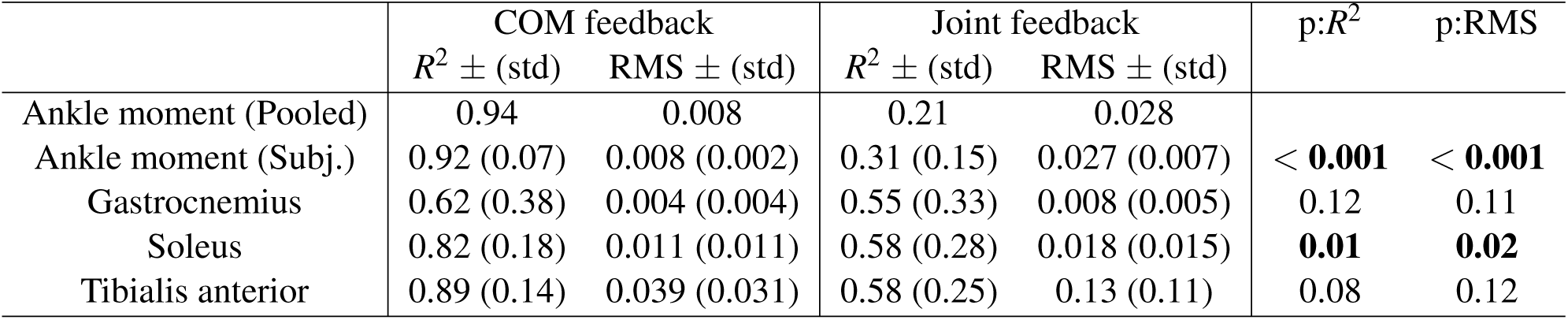
*Perturbed standing (data from [26])*: RMSE and *R*^2^ values of the reconstruction of the ankle joint moment and calf muscle activity in perturbed standing with COM feedback or ankle joint kinematics feedback. Joint moments are reported pooled over all subjects (Pooled) and for individual subjects (Subj.) with the standard deviation over subjects (std). P-values are reported of the paired ttest that compares the model with COM feedback and the model with ankle joint kinematics feedback. The results for muscle activity are only reported for individuals and not pooled over all subjects due to the limitations related to normalizing electromyography data.

**Figure 2:**
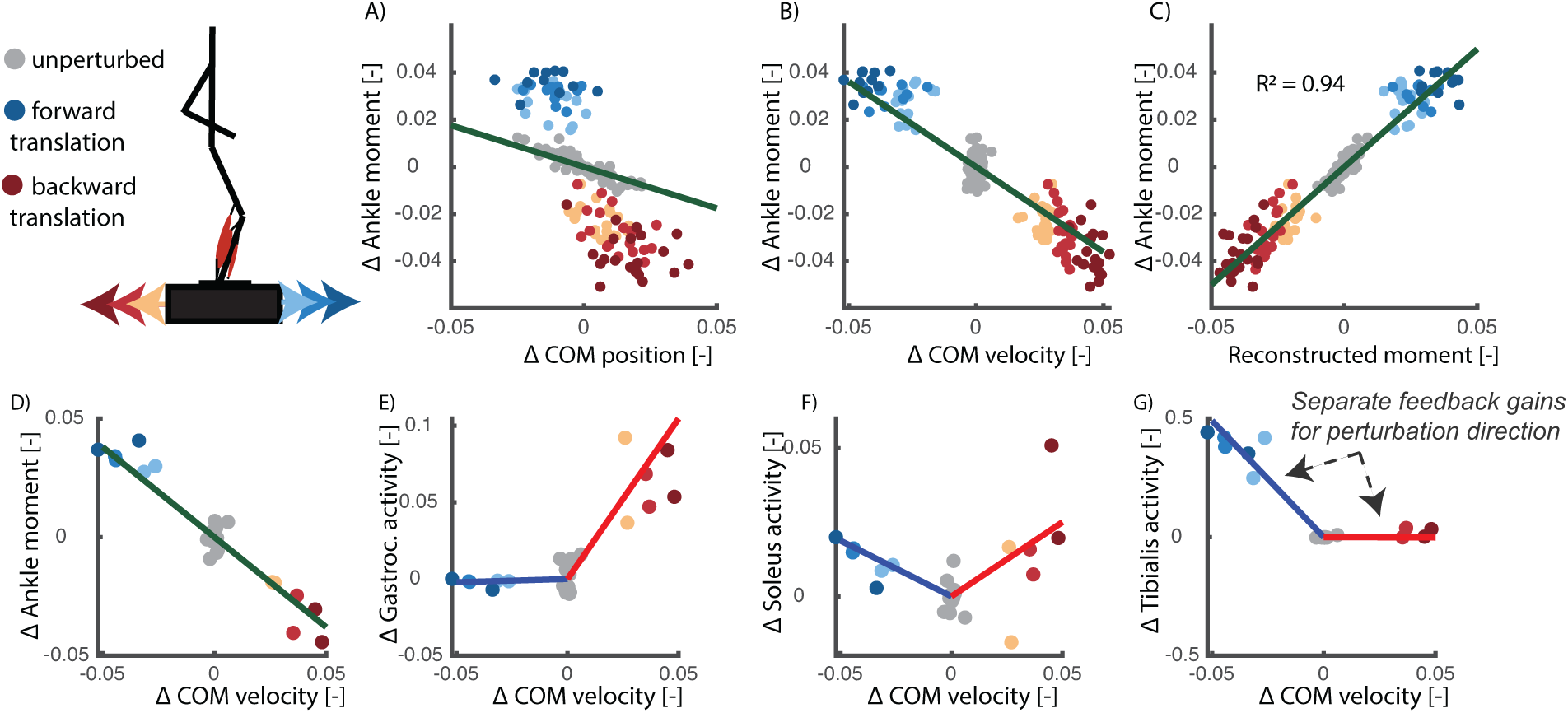
*Perturbed standing (data from [26])*: The relation between delayed COM kinematics and ankle joint moment and reactive muscle activity is visualised here at 150ms after perturbation onset. Each dot represents a single perturbation trial with a platform translation in forward (blue) or backward (red) direction with different magnitudes (color tint). A feedback model based on delayed COM position (A) and velocity (B) feedback can explain 94% of the variance in ankle moment (C) in response to support surface translations in forward and backward directions of different magnitudes. When analysing the data for each subject individually (D-G), we found that gastrocnemius and soleus activity increased with forward COM velocity and Tibialis anterior activity increased with backward COM velocity. Each dot in graphs (D-G) represents a single perturbation trial of a selected subject. Note that the electromyography data was analysed for individual subjects instead of pooled over all subjects due to the limitations related to normalizing electromyography data.

### 2.3 COM feedback explains ankle muscle activity and moment in perturbed walking

Similar as in standing balance, we found that reactive ankle joint moments and muscle activity can be explained by delayed feedback of COM kinematics, but not local joint kinematics feedback, in perturbed walking. COM kinematics explained ankle joint moments and muscle activity across perturbation protocols.

First, we analysed the relation between COM kinematics, joint angles and ankle moments in a dataset with pelvis-push perturbations [9] applied at toe-off of the contra-lateral leg. We found that the reactive ankle moment and muscle activity after the perturbation could largely be reconstructed by delayed COM feedback (Fig. 3, table 2). Tibialis anterior activity increased proportional to COM position and velocity in response to a backward directed perturbation (Fig. 4B) and gastrocnemius activity increased proportional to COM position and velocity in response to forward directed perturbations (Fig. 4C).

**Table 2:** *Pelvis push perturbations walking (data from [9])*: RMSE and *R*^2^ of the linear regression between delayed COM kinematics and reactive ankle muscle activity in perturbed walking. Joint moments are reported pooled over all subjects (Pooled) and for individual subjects (Subj.) with the standard deviation over subjects (std). P-values are reported of the paired ttest that compares the model with COM feedback and the model with ankle joint kinematics feedback. The results of muscle activity are only reported for individuals and not pooled over all subjects due to the limitations related to normalizing electromyography data.

| | COM-feedback | | Joint-feedback | | p-$R^2$ | p-RMS |
| --- | --- | --- | --- | --- | --- | --- |
| | $R^2 \pm \text{std}$ | RMS $\pm \text{std}$ | $R^2 \pm \text{std}$ | RMS $\pm \text{std}$ | | |
| **walking 0.62 m/s** |  |  |  |  |  |  |
| Ankle moment (Pooled) | 0.82 | 0.02 0.02 | 0.038 |  |  |  |
| Ankle moment (Subj.) | 0.84 (0.04) | 0.02 (0.01) | 0.19 (0.14) | 0.035 (0.004) | < **0.001** | < **0.001** |
| Gastrocnemius | 0.66 (0.19) | 0.06 (0.04) | 0.57 (0.18) | 0.07 (0.04) | 0.49 | 0.83 |
| Tibialis anterior | 0.75 (0.09) | 0.04 (0.01) | 0.50 (0.09) | 0.03 (0.01) | **0.002** | **0.004** |
| **walking 1.25 m/s** |  |  |  |  |  |  |
| Ankle moment (Pooled) | 0.71 | 0.02 | 0.20 | 0.027 |  |  |
| Ankle moment (Subj.) | 0.73 (0.08) | 0.02 (0.01) | 0.33 (0.08) | 0.024 (0.005) | < **0.001** | < **0.001** |
| Gastrocnemius | 0.66 (0.19) | 0.05 (0.02) | 0.56 (0.18) | 0.06 (0.02) | 0.15 | 0.19 |
| Tibialis anterior | 0.68 (0.10) | 0.03 (0.01) | 0.45 (0.20) | 0.04 (0.02) | **0.003** | **0.04** |

**Figure 3:**
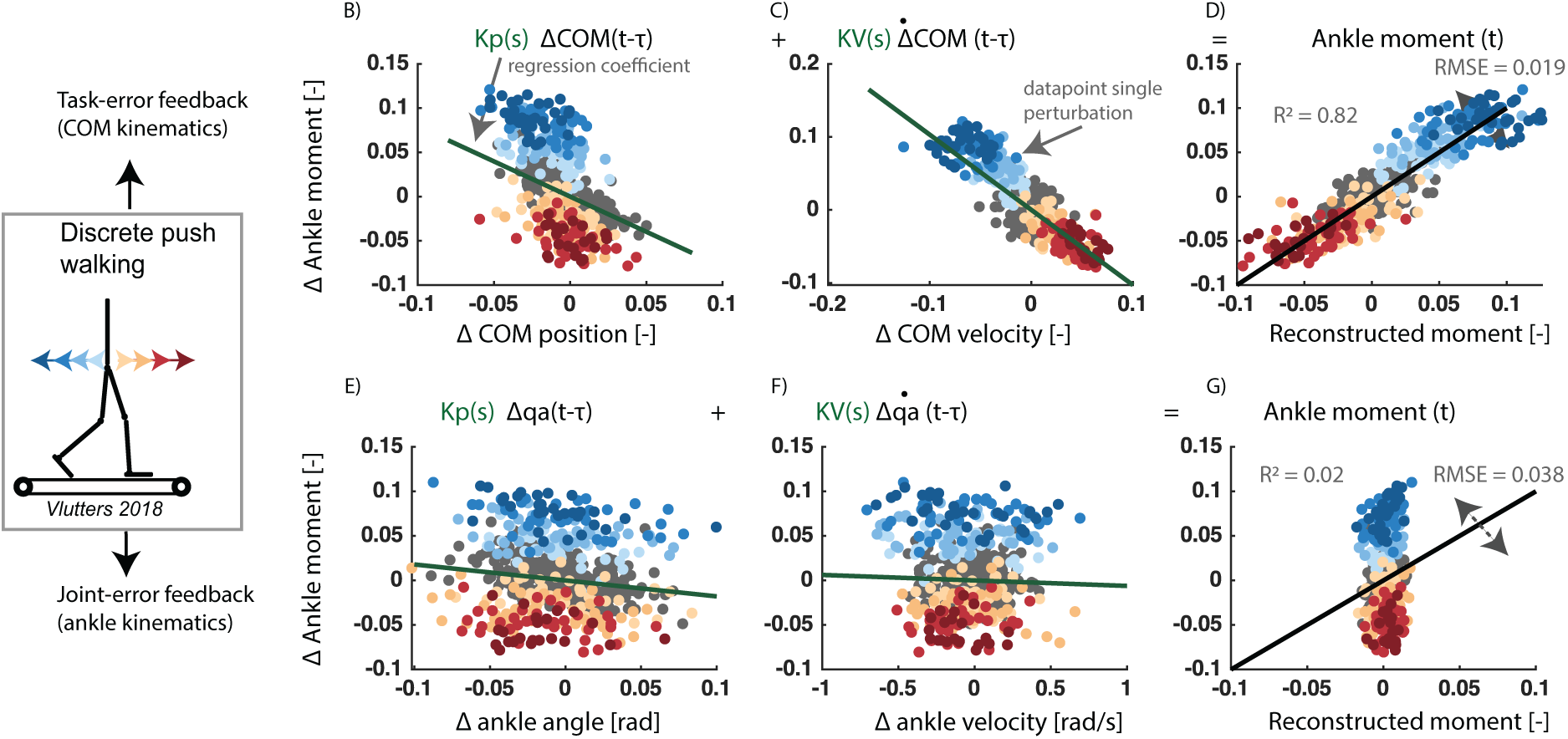
*Pelvis push perturbations walking (data from [9])*: Reactive ankle moment as a function of the deviation in COM position (A) and velocity (B) from the average reference trajectory during unperturbed walking (gray dots), in response to backward perturbations (blue dots) and forward perturbations (red dots). Least squares regression was used to estimate the position (*K*_*p*_) and velocity (*K*_*v*_) feedback gains, i.e. slope of green lines (A and B). The uncentered correlation coefficient (*R*^2^) and RMSE is represented in pane D by plotting the measured ankle moment as a function of the reconstructed moment based on COM kinematics. In the same dataset, a similar relation between reactive ankle moment and deviation in ankle joint angle (E) and velocity (F) was evaluated. Ankle moments reconstructed using COM kinematics (D) were more similar to the measured data than ankle moments reconstructed using ankle kinematics (G). This is data of walking at 0.6 m/s with the data pooled over all subjects.

**Figure 4:**
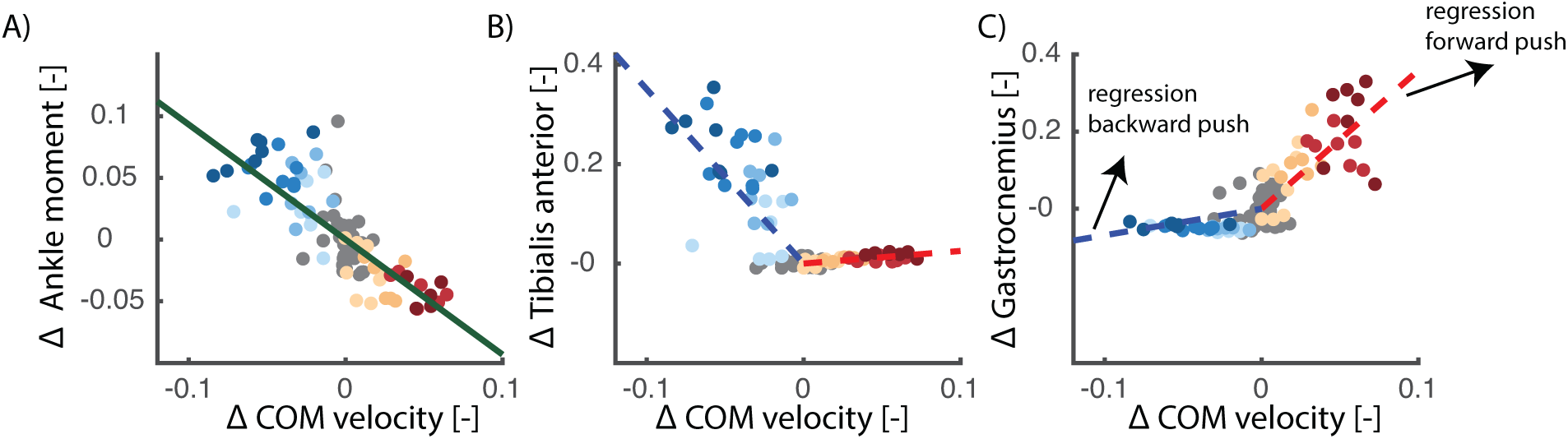
*Pelvis push perturbations walking (data from [9]):* Representative example of the relation between delayed center of mass velocity and reactive joint moments (A) and muscle activity (B and C) of one subject walking at 0.6 m/s with pelvis push perturbations. Unperturbed walking is visualised in gray, forward perturbations in red and backward perturbations in blue. There is a strong correlation between delayed COM velocity and the ankle moment (A). This relation is reflected in changes in tibialis anterior activity (B) and gastrocnemius activity (C). Tibialis anterior activity increases (quasi-linear) with negative COM velocity (i.e. blue regression line), and gastrocnemius activity increased with positive deviation in COM velocity (i.e. red regression line). Note that a representative example was selected rather than visualizing data pooled over all subjects, since this is only possible for the ankle moment and not for muscle activity due to the limitations related to normalization of electromyography data.

Second, we performed the same analysis in a dataset with a sudden increase or decrease in speed of the treadmill belts (also applied around toe-off of the contralateral leg). Similar as in the pelvis-push perturbations, we found that the reactive ankle moment after the perturbation could largely be reconstructed by delayed COM feedback (*R*^2^ = 0.81) (Fig. 5).

**Figure 5:**
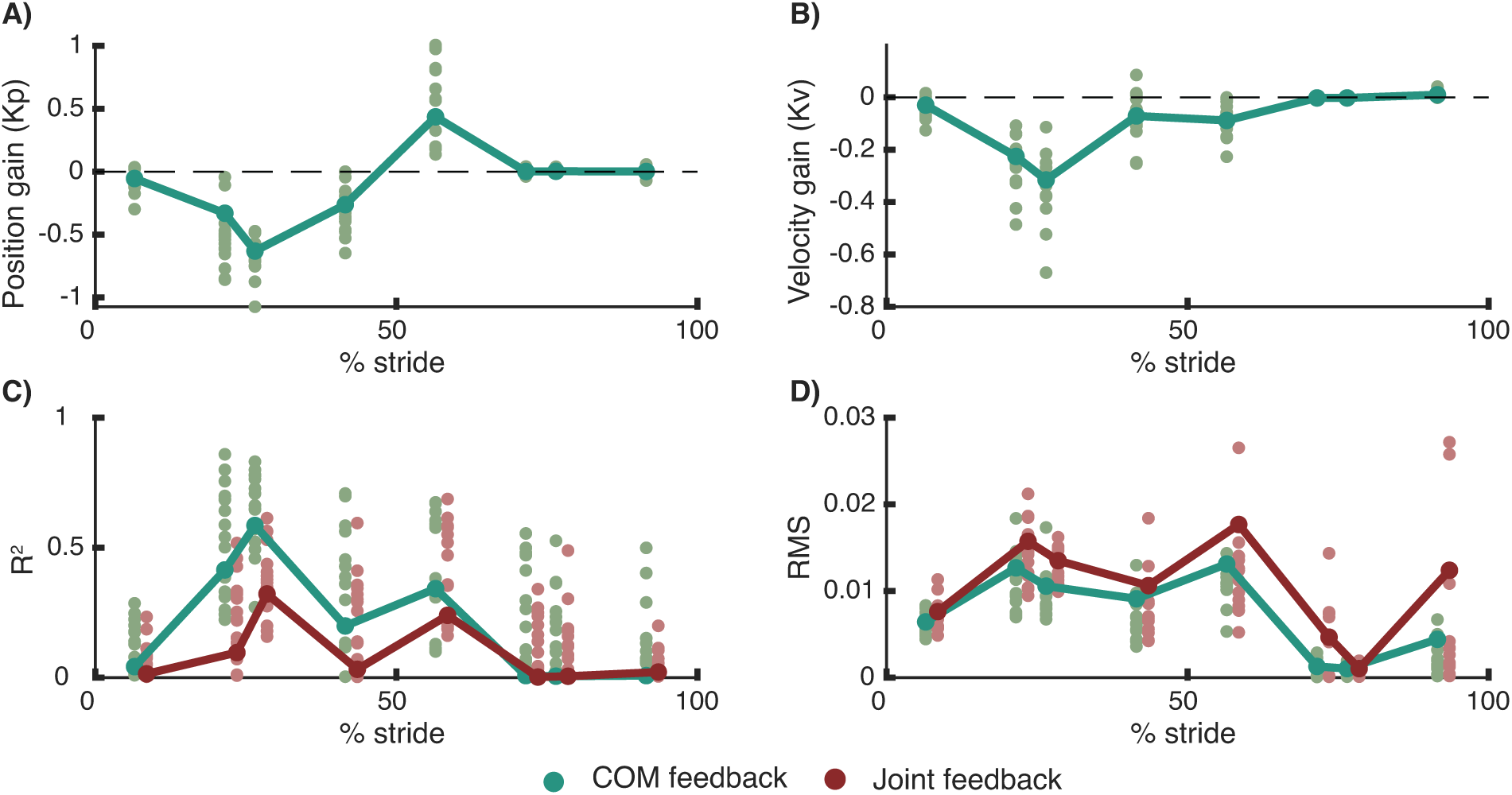
*Perturbed walking (data from [10])*: Gain modulation in response to beltspeed perturbations applied at different phases in the gait cycle. Reconstruction of the reactive ankle moments is better in a model with delayed COM feedback (green) compared to local joint level feedback (R) (C,D). The proportional feedback of COM position (A) and COM velocity (B) is modulated during the gait cycle. The bars and large dots represent feedback gains estimated based on the pooled data over all subjects. The small dots represent gains estimation in individual subjects. The perturbation onset timing in the gait cycle had a significant influence on the position gains, velocity gains, *R*^2^ and RMS values.

In both experiments, ankle moments reconstructed using COM kinematics were more similar to the measured data than ankle moments reconstructed using ankle kinematics (table 2, Fig. 5).

### 2.4 Modulation COM feedback during the gait cycle and with walking speed

We found that the COM feedback gains are dependent on the phase in the gait cycle and are modulated with walking speed both when perturbations happen at discrete time instants and continuously. The observed change in feedback gains in the gait cycle in combination with the high *R*^2^ values during midstance and low *R*^2^ values during initial and terminal stance (Fig. 5) indicates that the control of COM kinematics (1) changes during the gait cycles, (2) is most pronounced during mid-stance and (3) is only active when the foot is in contact with ground.

First, we used a dataset with changes in the belt speed in anterior and posterior direction at discrete instances in the gait cycle [10]. At the level of ankle joint moments, estimated feedback gains (*K*_*p*_ and *K*_*v*_) depended on the phase in the gait cycle (one representative subject: figure 6A, all subjects figure: 5). The position and velocity gains are highest during mid-stance and lowest during the swing phase. The variance in ankle moment explained by the COM feedback model (*R*^2^ values) is highest during mid-stance (Fig. 5). A similar modulation is observed at the level of reactive ankle muscle activity (Fig. 6). The proportional increase in Gastrocnemius and Soleus muscle activity with deviations in COM kinematics in response to forward perturbations was highest during mid-stance (Fig. 6). The proportional increase in tibialis anterior activity with deviations in COM kinematics in response to backward perturbations was most pronounced during the first half of the stance phase. Remarkably, during mid-stance, calf muscle activity was also inhibited proportional to backward displacement and velocity of the COM in backward directed perturbations (Fig. 6). This indicates that both muscle excitation and inhibition are proportional to deviations in COM kinematics in this phase of the gait cycle.

**Figure 6:**
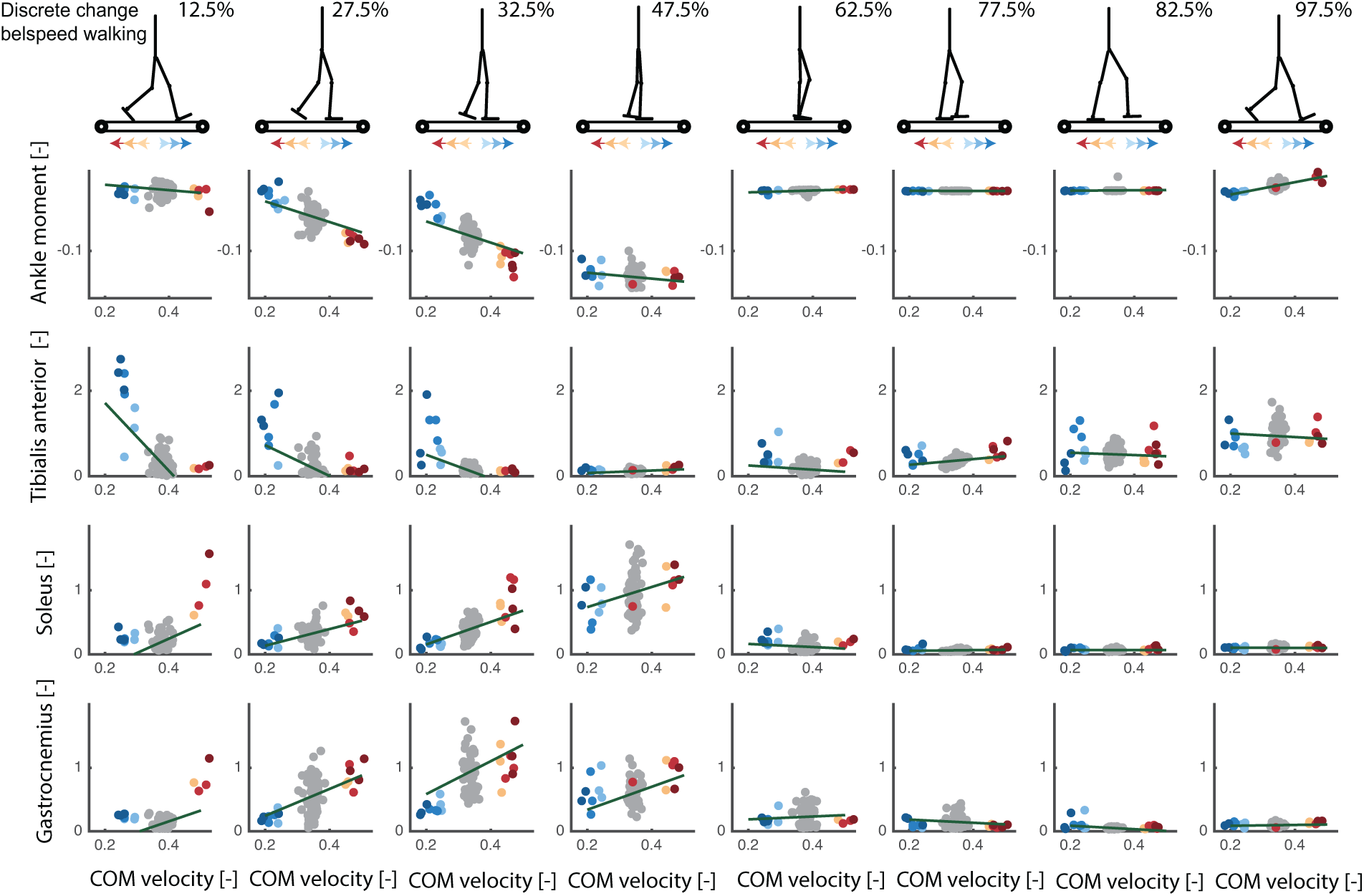
*Discrete treadmill perturbations walking (data from [17]*): Representative example of the relation between deviations in delayed center of mass velocity and reactive joint moments (row 1) and activity of the tibialis anterior (row 2), soleus (row 3) and gastrocnemius (row 4) in response to perturbations during walking. Perturbations were applied at four instances during the stance phase of the left leg resulting in eight responses at joint level when combining data of the left and right leg (e.g. right leg is in swing during mid-stance of the left leg). Unperturbed walking is visualised in gray, increases in belt speed (i.e. forward fall) in red and decrease in belt speed in blue (i.e. backward fall).

Second, we used a dataset with continuous changes in the speed of both belts while walking at 0.8, 1.2, and 1.6 m/s [27]. The main advantage of this data is that approximately 400 strides of perturbed walking could be analysed for each subject at multiple walking speed. To investigate phase-dependency of the reactive ankle moment within the stance phase, we divided the stance phase in 16 bins and computed a linear model for delayed COM feedback to ankle joint moment in each of those bins (Fig. 7). Similar as in the discrete perturbations, we found that the changes in ankle joint moments during the gait cycle are closely related to COM position and velocity (*R*^2^ up to 0.7 during mid-stance, figure 8). Similar to the discrete perturbations, the *R*^2^ values and feedback gains are low during early and late stance and high during mid-stance (Fig. 8). In addition, we found that feedback gains and the variance explained (*R*^2^) by the linear regression decreased with increasing walking speed (Fig. 8). Note that also for this dataset, the reconstructed ankle moment is more similar to the measured data when using COM feedback than using joint kinematics feedback for walking at slow and medium speeds(p-*R*^2^ *<* 0.001 and p-*RMS <* 0.001 for walking at 0.8 *m/s* and 1.2 *m/s*) but not for walking at faster speeds (p-*R*^2^ = 0.71 and p-*RMS* = 0.59). The average *R*^2^ and RMS values during the stance phase are respectively 0.38 and 0.014 with COM feedback and 0.23 and 0.018 with ankle kinematics feedback. The feedback model was not evaluated at muscle level since no electromyography data was collected in this dataset.

**Figure 7:**
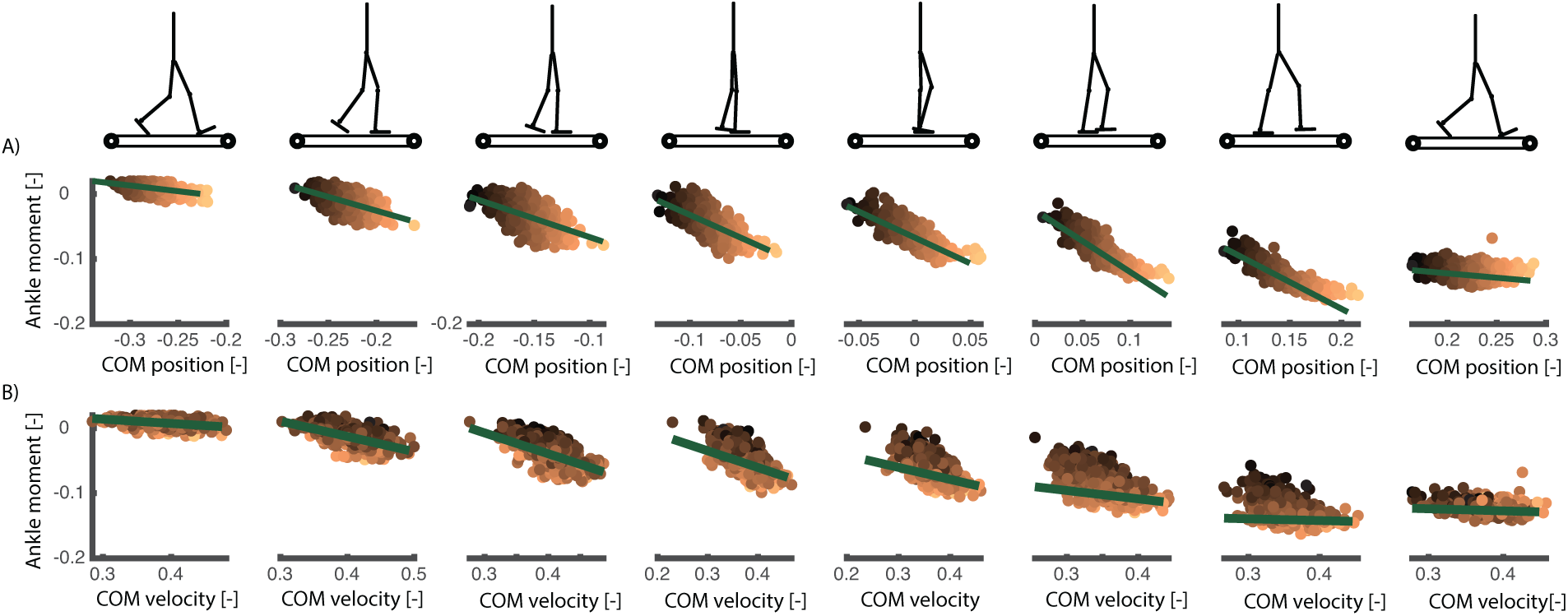
*Continuous treadmill perturbations walking (data from [27]*) : Representative example of the relation between ankle joint moment and COM position and velocity in walking with continuous changes in the speed of both belts. The different columns in the plots are time bins equally spaced during the stance phase of walking. The colored dots represent the different strides of this subjects (with the color tint representing the deviation in COM position), the slopes of the green green lines are the resulting position and velocity gains from the least squares regressions. The change in slope of the green lines show the phase-dependency of the linear regression between COM position and velocity and the ankle moment during perturbed walking.

**Figure 8:**
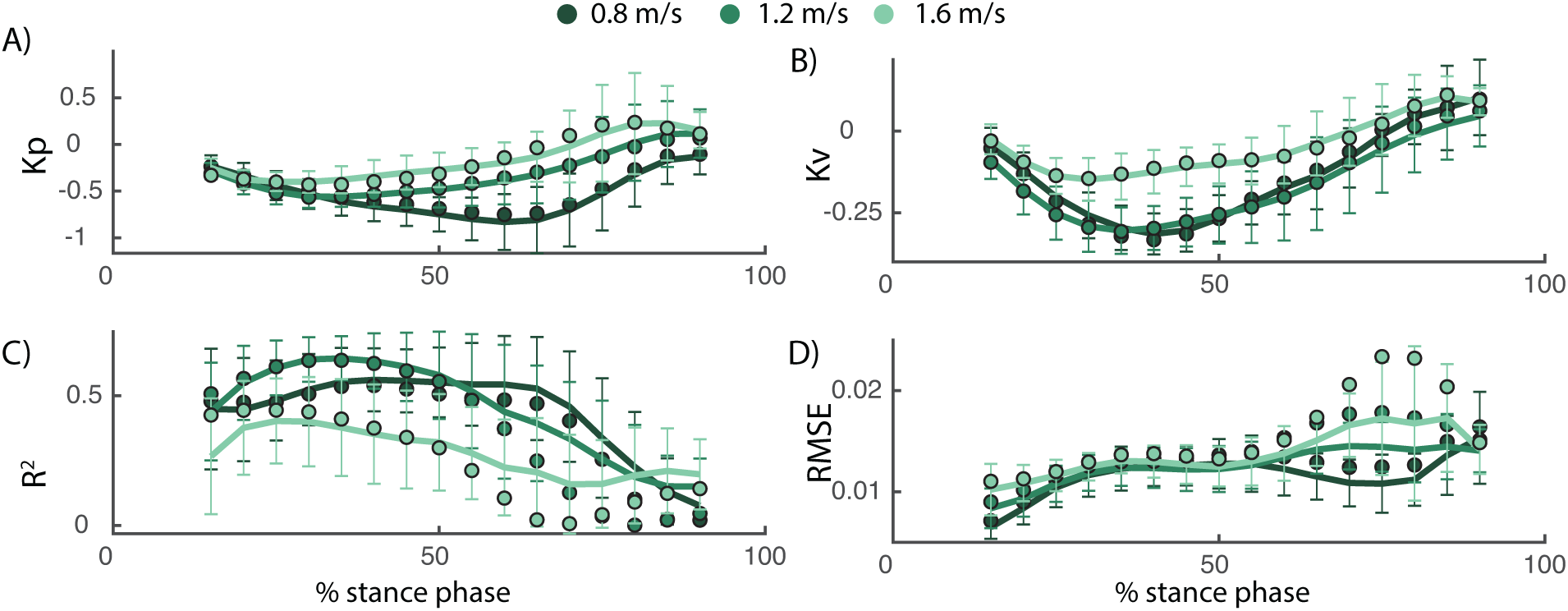
*Continuous treadmill perturbations walking (data from [27]*): The COM position (A) and velocity (B) feedback gains describing the reactive ankle joint moment depend on the walking speed. Both the position and velocity gains decrease with increasing walking speed. *R*^2^ values decrease and RMSE increase with increasing walking speed, indicating a worse fit of the feedback model as walking speed increases. The errorbars and lines represent the average and standard deviation of the fit for individual subjects, the dots represent the results from the analysis based on pooled data over all subjects.

To evaluate if the modulation of feedback gains is important to describe the ankle strategy during the gait cycle, we compared the model with phase-dependent gains with a model with constant gains during the gait cycle (i.e. *K*_*p*_ and *K*_*v*_ are constant 1). For this analysis, we used the dataset with continuous treadmill perturbations during walking as this dataset contained perturbation data of many steps throughout the entire gait cycle. Due to the lower number of variables, we found that the *RMSE* increased in the model with constant gains compared to the model with variable gains, especially at the end of the stance phase (Fig. 9). Similarly, the *R*^2^ value decreased in the model with constant gains, especially during the first part of the stance phase. However, the average decrease in *R*^2^ values (0.08) and in RMSE (0.002) in the constant gain model compared to the phase dependent gain model was small (Fig. 9).

**Figure 9:**
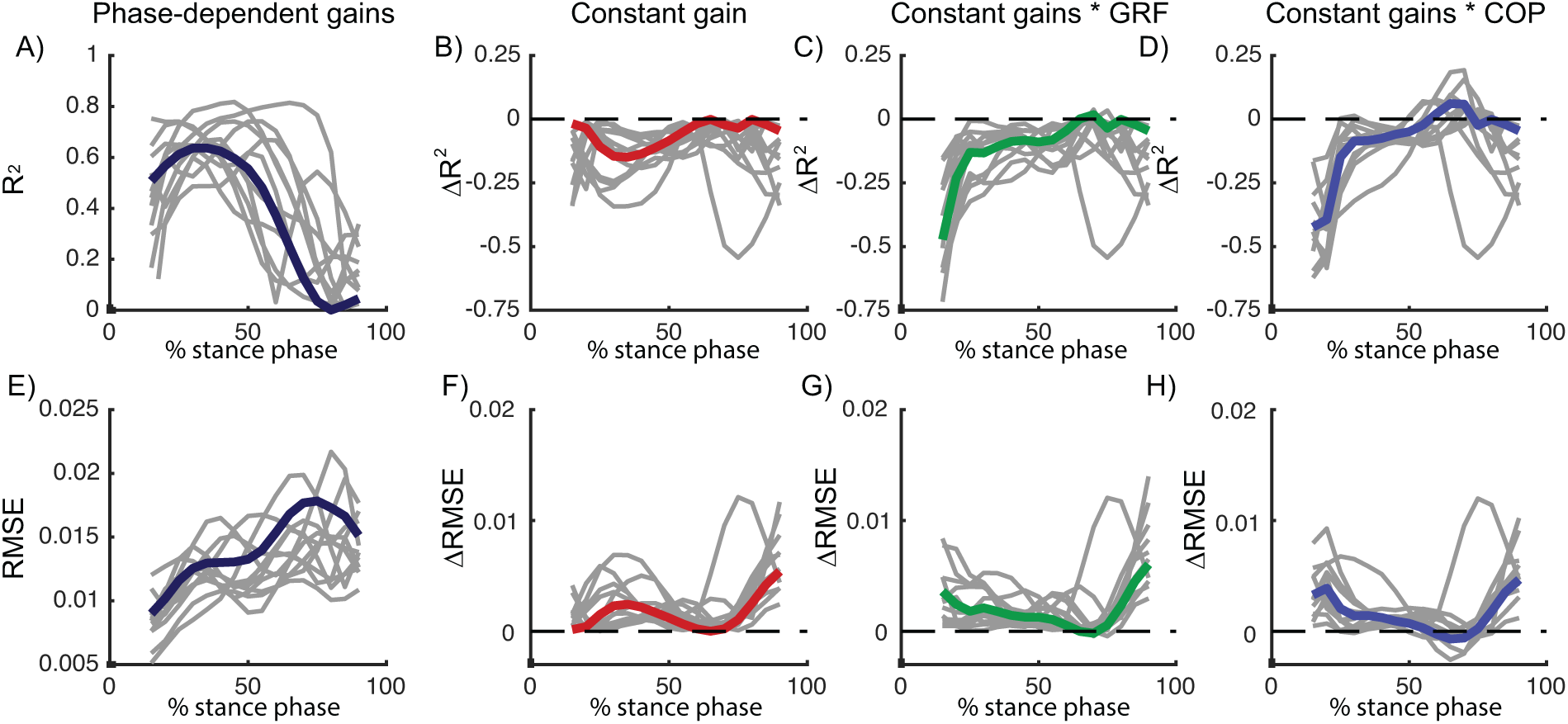
*Continuous treadmill perturbations walking (data from [27])*: *R*^2^ and RMSE of the linear regression between COM kinematics and ankle moment in a model with phase-dependent feedback gains (A, dark blue). Relative to the model with phase-dependent gains, *R*^2^ and RMSE values increased in models with constant feedback gains (B, red), constant feedback gains multiplied by vertical ground reaction force (C, green) and constant feedback gains multiplied by the COP position (D, blue). Data pooled over all subjects are visualised in color and results for individual subjects are visualised in gray. A similar increase in RMSE and decrease in *R*^2^ was found in the three models, indicating that the quality of fit did not increase with information from the vertical force or COP. Results of the statistical analysis can be found in table 3

### 2.5 Modulation COM feedback during gait cycle cannot be fully explained by tactile information from the foot

We hypothesized that the observed modulation of feedback gains during the gait cycle might be driven by cutaneous information from the foot. To test this hypothesis, we modified the constant gain model to include delayed feedback from cutaneous sensory information represented by either the location of the COP in the foot or the vertical ground reaction force. To evaluate whether this model could explain the phase-dependent modulation of COM feedback, we used both a model in which the constant COM feedback gains were multiplied by delayed feedback from the vertical ground reaction force (*F*_*y*_) or COP location (*COP*_*b*_, minimal distance between the COP and the bound of the foot).

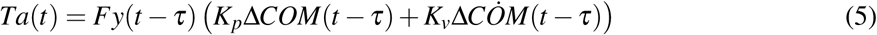

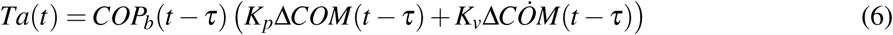

Modulation of feedback gains by the vertical ground reaction force or COP did not improve the fit (*R*^2^ and RMSE) compared to the model with constant gains (Fig. 9). In addition, for walking at 1.2 m/s, the modulation of constant feedback gains with the vertical ground reaction force resulted even in a significant decrease in the *R*^2^ values compared to the model with constant gains (p = 0.01, see table 3 for details).

**Table 3:**
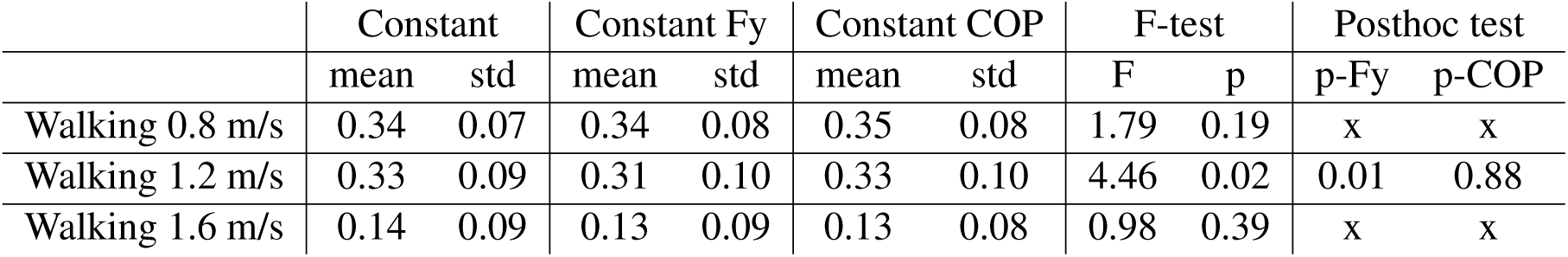
Statistical comparison of the *R*^2^ values, averaged over the stance phase, of models with (1) constant feedback gains (*Constant*), (2) constant feedback gains modulated by the vertical ground reaction force (*Constant Fy*) and (3) constant feedback gains modulated by the center of pressure position (*Constant COP*).we found only for walking at 1.2 m/s a significant difference between the three models (repeated measures anova). For this walking speed, only the modulation of constant feedback gains with the vertical ground reaction force resulted in an significant increase in the *R*^2^ values compared to the model with constant gains (post-hoc testing with bonferroni correction). Note that the post-hoc test was not evaluated when the F-test was not significant (x in the table).

## 3 Discussion

### 3.1 Common sensorimotor transformations during standing and walking

We showed that common sensorimotor transformations can explain the ankle response in perturbed standing and walking. Both reactive ankle muscle activity and joint moments can be reconstructed by a linear combination of delayed COM position and velocity in perturbed standing and walking, rather than with local ankle joint kinematics feedback. Similar as in standing balance [3], we showed that the reactive EMG activity and ankle moments was better described by delayed feedback of COM kinematics across walking speeds and different perturbation modalities, compared to delayed feedback of ankle kinematics. This result supports the hypothesis that the nervous system estimates relevant task-level variables from multi-sensory information to activate muscles to control balance in response to perturbations [6, 18]. The neural mechanisms behind the task-level feedback of COM kinematics remains however unclear. Sensory perturbation experiments, with for example galvanic stimulation, might provide more insight in neural integration of sensory information from multiple sources to estimate COM kinematics.

### 3.2 Modulation feedback control during walking

We hypothesized that task-level feedback from COM kinematics are modulated during walking to compensate for changes in body configuration during the gait cycle. We indeed found that estimated feedback gains changed over the gait cycle in two datasets with perturbations during different phases of the gait cycle [10, 27]. Both the feedback gains and the variance in ankle moment explained by the linear regression were high during mid-stance but low during initial and terminal stance and during swing. Hence, the ankle strategy is mainly used during mid stance, which is logical from a mechanical point of view. In the ankle strategy, balance is controlled by modulating the interaction with the ground [8]. More specifically, activity of the ankle muscles changes the COP position in the foot to control balance [8, 10]. This mechanism has a high potential to control COM movement during mid-stance since the COP is approximately in the middle of the foot and can move in forward and backward directions. Hence, our results indicate that the proportional feedback of COM kinematics is modulated to exploit this change in potential to adjust the COP position within the foot.

Our results suggest that the modulation of COM feedback gains is similar to the modulation of local reflexes during walking. The similarity in phase-dependency and speed-dependency of COM feedback gains and H-reflex during the gait cycle suggests that modulation of COM gains and local reflexes are related. Both the H-reflex of the soleus and gastrocnemius and the gains of the COM feedback that explains the ankle strategy after perturbations are high during mid-stance [22] and low during early stance, terminal stance and during the swing phase. Furthermore, we also observed that the gains of the COM feedback decrease with increased walking speed and are higher in standing compared to walking, which is similar to the modulation of h-reflex magnitude with walking speed [23].

The sensory mechanism underlying the observed modulation in COM feedback gains during the gait cycle remains unclear. Given changes in foot-ground interaction throughout the gait cycle and the dependency of reflex activity on tactile information from the foot during walking [24, 25], we hypothesized that information from the COP location and vertical ground reaction force could explain the modulation of reflex gains. In contrast with this hypothesis, we found that the fit of the feedback model did not improve when scaling the feedback gains with COP information or the vertical ground reaction force (figure 9). We believe that there are two potential explanations for this observation. First, this might indicate that tactile information in the foot cannot explain the modulation of COM feedback gains during the gait cycle. Second, this might also indicate that the simplifying assumption of modelling information from cutaneous sensors with linear feedback of vertical ground reaction force or COP location is not valid. In the future, cutaneous stimulations experiments could generate more insight in the modulation of task-level feedback gains during walking [25].

The decrease in *R*^2^ values and feedback gains with increased walking speed indicates that task-level feedback of COM kinematics is less important in fast walking. There are multiple explanations for the decreased importance of task-level feedback with increased walking speed. These changes in task-level feedback might reflect an increase in inherent stability of the skeleton system with walking speed. Alternatively, subjects might rely less on an ankle strategy and more on a stepping strategy at higher walking speeds. The ankle strategy can act sooner than the stepping strategy, which has no effect before the next foot pcontact, might be less important during fast walking [12]. Step frequency increases with increased walking speed, thereby reducing the duration of the swing phase. As a result, the next foot placement occurs earlier and therefore the delay between perturbation onset and foot placement to control balance decreases. In addition, the shorter duration of the stance phase with increased walking speed might limit the potential of the ankle strategy to control balance.

When comparing the different types of perturbations in walking (Fig. 10), we found that both the position and velocity feedback gains are higher in the pelvis push perturbations (walking at 1.25 m/s) compared to the support surface perturbations (walking at 1.2 m/s, figure 10). This might be the result of oversimplification of task-level feedback with linear feedback of COM kinematics. We could only explain up to 80 % of the variance in ankle moment with this simple feedback model. Non-linear feedback or feedback from other task-level variables such as COM accelerations or head orientation [6] might contribute to reactive muscle activity and joint moments in perturbed walking.

**Figure 10:**
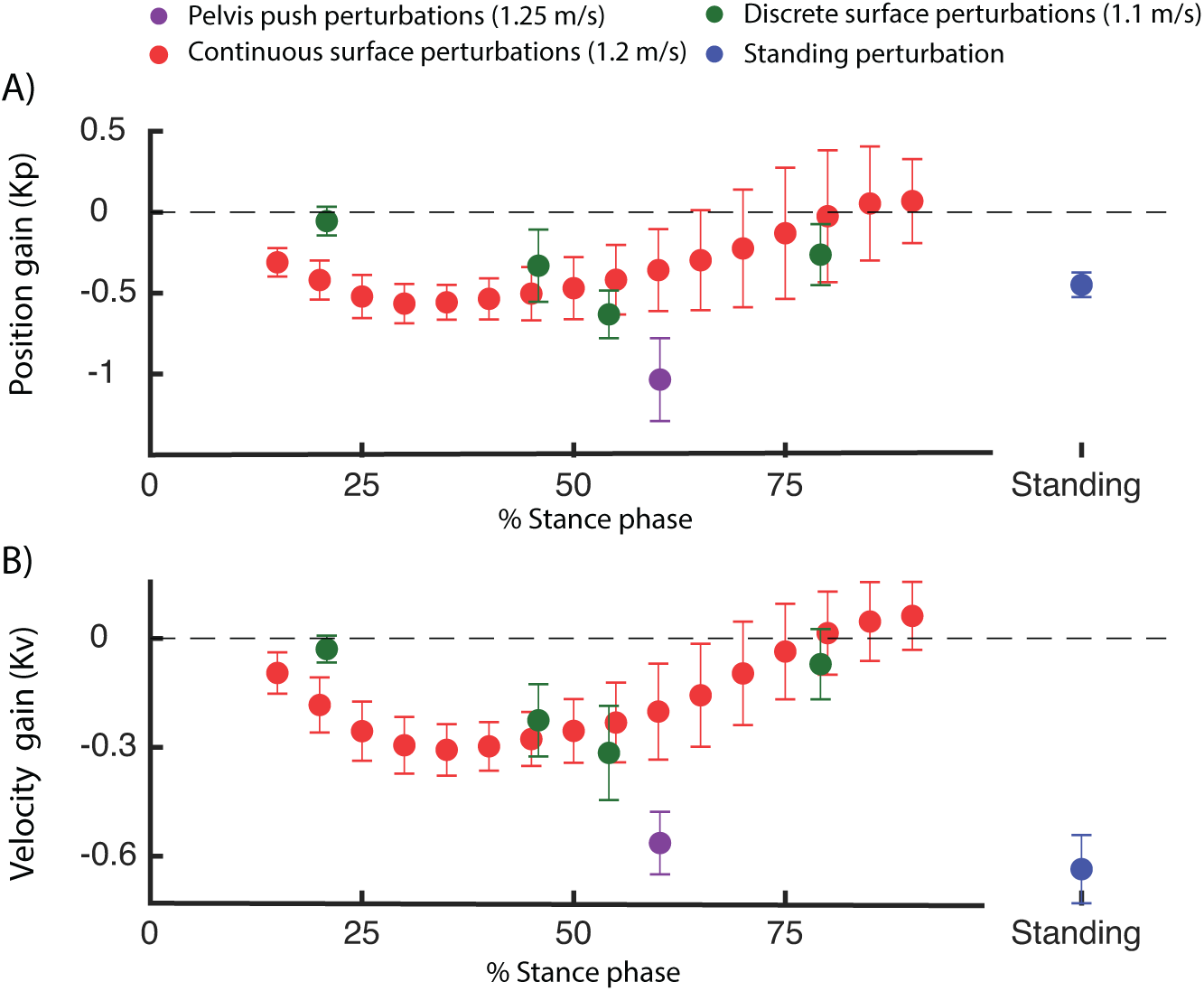
Position and velocity gains computed in the dataset with perturbed standing (green [26]), walking with discrete (blue [10]) and continuous(red [27]) surface perturbations and discrete pelvis push perturbations (purple [9]). The feedback gains for walking are presented as a function of the percentage of the stance phase. Note that the percentage in the stance phase represents the phase at which the feedback model was evaluated (i.e. 250 ms after perturbation onset).

**Figure 11:**
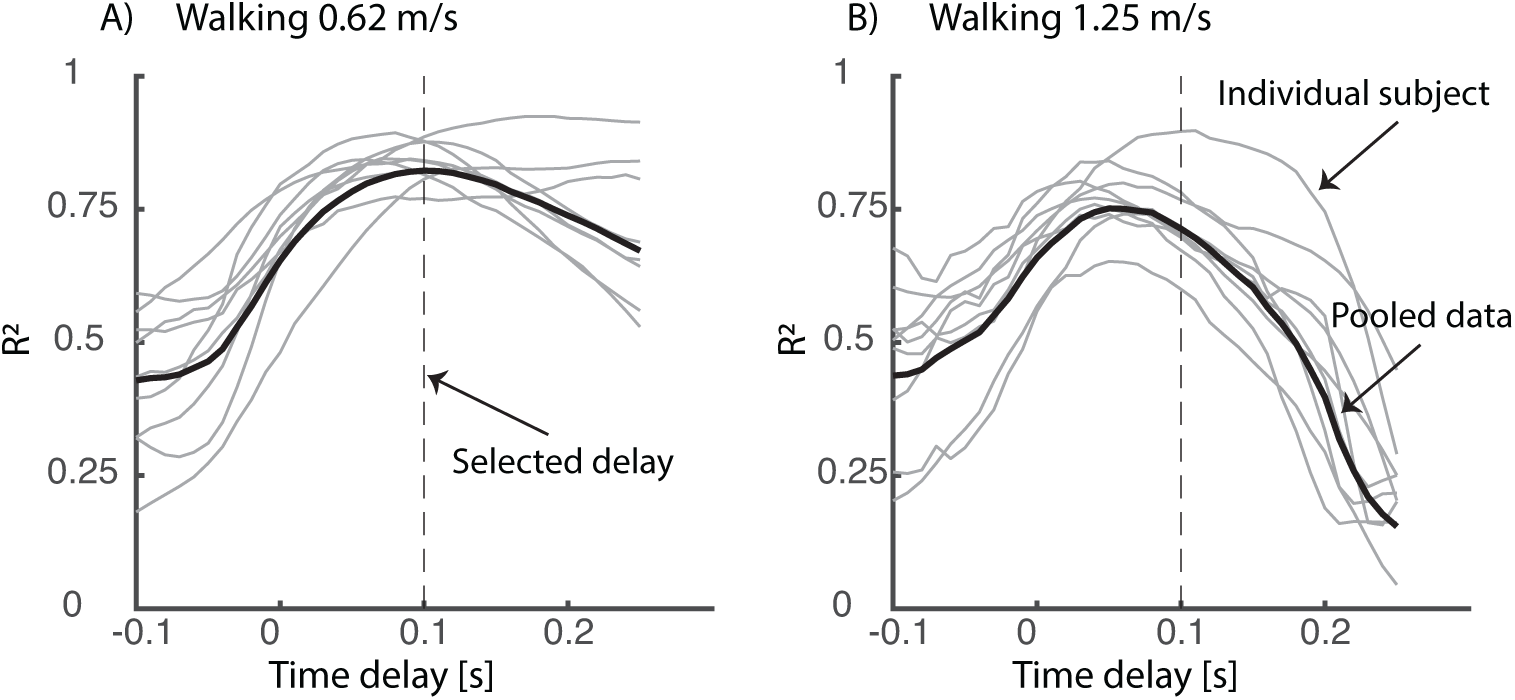
*Sensitivity of the time delay:* The variance explained by the model is influenced by the neural time delay (*τ*_*T*_ in equation 1). We performed this analysis for the relation between reactive ankle moment and COM kinematics in the pelvis push perturbations [9]. Note that time delay in this study was selected based on literature [29] and was not based on this sensitivity analysis.

**Figure 12:**
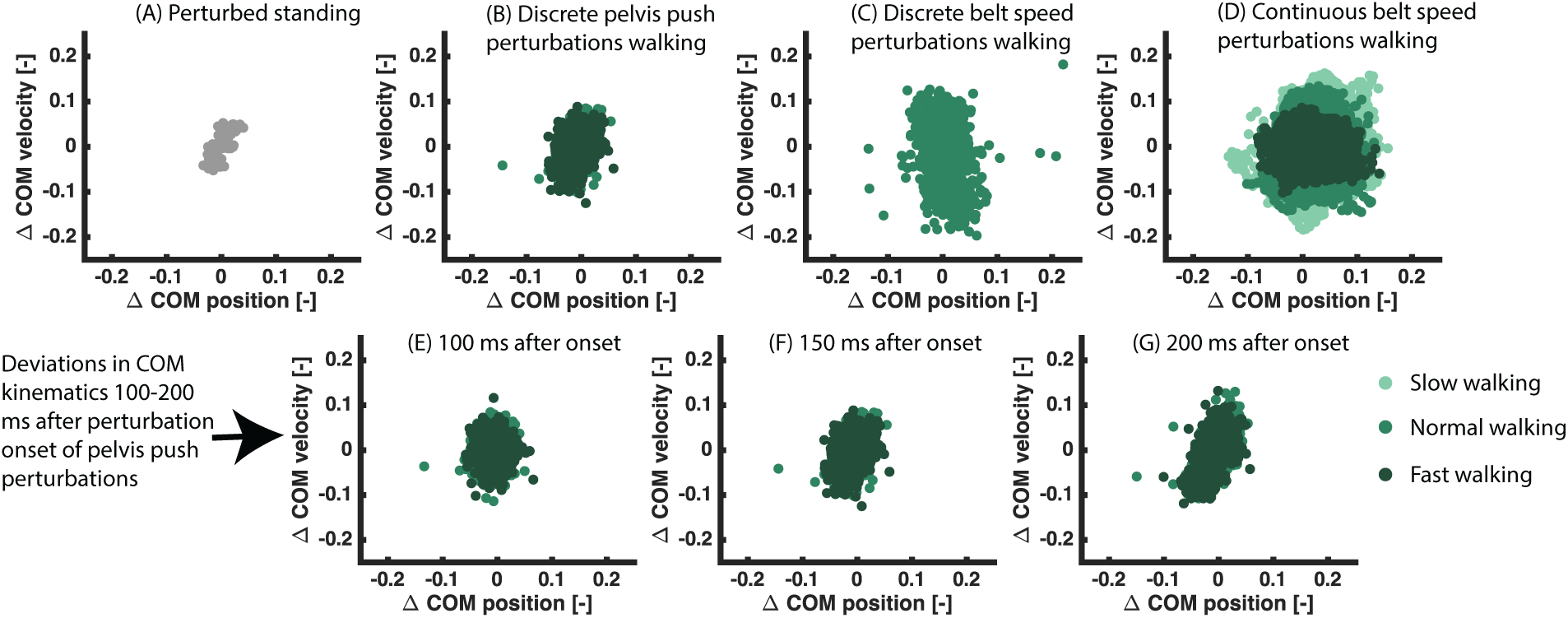
*Deviations in COM position and velocity in the different types of perturbations:* The deviation in COM position and velocity is similar in the different datasets of perturbed walking (B-D) and is larger in perturbed walking compared to perturbed standing (A). A similar deviation in COM kinematics is observed when analysing the data 100ms (E), 150ms (F) and 200ms (G) after perturbation onset in the pelvis push perturbations.

### 3.3 Contribution of reactive muscle activity

The relation between COM kinematics and reactive muscle activity shows that the reactive ankle joint moments do not solely result from visco-elastic properties of active muscles, but result from active feedback control (Fig. 4). Based on the relation between joint moments and delayed COM kinematics, it is unclear if the modulation of ankle moment is indeed an active control mechanism. Given that a similar correlation was observed at the level of reactive activity of the calf muscles confirmed the important contribution of active feedback control.

Despite the clear contribution of active control mechanisms, it is hard to identify the relative importance of active feedback control and passive dynamic contributions to the ankle moment. Musculoskeletal modelling could be used to identify muscle stiffness and damping using muscle models [30]. This would enable estimation of joint stiffness and damping. However, this relies on accurate methods to estimate the muscle state from motion capture data (e.g. [31]) and on accurate models of muscle contraction dynamics [32].

### 3.4 Derive neuromechanical models for balance control during walking

Our findings have important implications for modelling neuromechanics of walking. In current models, balance is mainly controlled by foot placement [15,33]. Here, we showed that subjects also adjust the ankle joint moment in response to perturbation during walking and that this strategy can be modelled using delayed COM position and velocity feedback. Hence, we propose an additional supra-spinal feedback loop to model the ankle strategy during walking, similar to the neuromechanical models of perturbed standing [29]. Since this additional feedback can describe largely the ankle joint moment in response to different types of balance perturbation across walking speeds, we believe this is an important extension of the current state-of-the art neuromechanical models. We believe that this especially useful in the control of wearable robotic devices. Current neuromechanical models that are used to control exoskeletons and prosthesis [34] lack balance control mechanisms for the ankle joint. We hypothesize that this additional feedback loop will facilitate shared control of balance when using neuromechanical models in the control of wearable robotic devices.

## 4 Materials and methods

We used existing data from multiple perturbation experiments in healthy young and older subjects to derive sensorimotor transformations for the ankle strategy during standing and walking (see figure 1 for an overview).

### 4.1 Experimental methods

Integrated 3D motion capture was used in four datasets to measure the human response to perturbations during standing and walking. An overview of the different datasets and measurements is given in figure 1.

- Standing balance was perturbed using support surface translations triggered at a discrete time instance. Integrated motion capture data was collected in 8 young (age 21*±* 2std) and 10 older adults (age 67 *±* 3std). The perturbation protocol and data collection is extensively described in [26]. In short, standing balance was perturbed by means of a sudden forward and backward platform translation with respectively fast, medium and slow acceleration profiles (Fig. 1), as well as medio-lateral platform translations, but only forward and backward perturbations were analyzed in this study. Perturbations were applied in a random order. Whole body motion was recorded using 3D motion capture with an extended plug in gait marker (Vicon, Oxford Metrics). Ground reaction forces were collected from two AMTI force plates (AMI, Watertown, USA) embedded in the platform. This dataset has already been used to analyse kinematic strategies in response to backward surface translations in [17]. Here, we additionally analyzed the response to forward support surface translations and reactive muscle activity of the tibialis anterior, soleus and gastrocnemius, which was collected using surface electromyography (Cometa).
- Steady state walking was perturbed by means of a sudden change in speed of the treadmill belt, triggered at specific phases of the gait cycle. Integrated motion capture data was collected in 18 young (age 21 *±* 2 std years) and ten older adults (age 71*±* 4 years). The perturbation protocol and data collection in extensively described in [10]. In short, the perturbations consisted of a sudden increase or decrease in speed of the treadmill belt with two different magnitudes. These perturbations were applied at four different phases of the gait cycle. Medio-lateral treadmill translations were also part of the perturbation protocol but were not analysed in this study. All perturbations were applied in a random order. All subjects walked at 1.1 m/s on a split-belt treadmill. Whole body motion was recorded using 3D motion capture with an extended plug in gait marker set (Vicon, Oxford Metrics). Ground reaction forces were collected on a split-belt treadmill (Motek-ForceLink). Muscle activity of the gastrocnemius, soleus, tibialis anterior was measured using surface electromyography (Bortec Octopus 8 channel electromyography).
- Steady state walking was also perturbed by means of an external force (push) at the pelvis. Integrated motion capture data was collected in 9 subjects (age 25*±* 2 std years). The perturbation protocol and data collection is extensively described in [9]. In short, the perturbations consisted of a forward of backward directed push at the pelvis with four different magnitudes. Forward and backward push perturbations were applied in a random while the subjects walked at 0.62 m/s and 1.25 m/s. Whole body motion was recorded using 3D motion capture with an full-body marker protocol (VisualEyez-II). Ground reaction forces were collected on a split-belt treadmill (MotekForceLink). Muscle activity of the tibialis anterior and gastrocnemius was measured using electromyography (Delsys).
- Steady state walking was also perturbed by means of continuous changes in the speed of the treadmill belts. Integrated motion capture data was collected in 15 young subjects (age 24*±* 4 std years). The perturbation protocol and data collection is extensively described in [27]. In short, balance was perturbed continuously using three pseudo-random belt speed control signals, with mean velocities of 0.8 *m/s*, 1.2 *m/s* and 1.6 *m/s*. Each trial with continuous perturbations consists of 480 seconds walking, resulting in approximately 500 gait cycles of perturbed walking for each subject and each walking speed. Whole body motion was recorded using 3D motion capture with an full-body marker protocol. Ground reaction forces were collected on a split-belt treadmill (MotekForceLink). No electromyography data was collected in this study.

### 4.2 Inverse kinematic and dynamic analysis of the experimental data

Joint kinematics and kinetics were computed using a scaled generic musculoskeletal model with 23 degrees of freedom (gait 2392) in OpenSim [35]. This model was scaled to the anthropometry and mass of the subject based on the marker positions and ground reaction forces in the static trials. Joint kinematics of the scaled model were computed from the recorded marker trajectories using a Kalman smoothing algorithm [36]. Joint kinetics were computed based on the equations of motion of the model with OpenSim’s inverse dynamics tool. The motion capture dataset with continuous perturbations was processed using the open source software to compute joint kinematics and kinetics provided with the manuscript [27]. Whole body COM position and velocity was computed from the joint kinematics and the model of the skeleton (i.e. kinematic approach), and was expressed relative to the position and velocity of the base of support (i.e. foot in contact with the ground).

### 4.3 normalization of data

Joint kinematics, kinetics and muscle activity data were normalized to facilitate comparison between subjects and datasets to enable estimation of feedback gains based on data pooled over all subjects. Similar as in [28], COM positions were normalized by *l*_*max*_, speeds by 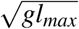, torques by *mgl*_*max*_ and muscle activity by MVC values for the standing data and by the peak average activity observed during unperturbed walking for the walking data.

### 4.4 Linear regression

We used least squares regression to obtain linear models between the deviations in ankle and COM kinematics (inputs) and reactive ankle joint moment and muscle activity (outputs) after perturbation. We first computed the mean of the inputs and outputs in the unperturbed standing and walking data. Subsequently, we computed the deviation from the means in the perturbed standing and walking data. For example, in standing balance, we first used the data at 0.5 seconds before perturbation onset, averaged over the multiple perturbation trials, to determine the mean of the unperturbed COM kinematics, joint moments and muscle activity. We then computed the deviation from this mean in each perturbed and unperturbed trial. These deviations were used as input for the least squares regression. The same method was used in perturbed walking, but the inputs and outputs were expressed as a percentage of the stance and swing phase. The linear regression was evaluated at 150ms after perturbation onset in all datasets because a large deviation in COM kinematics was observed in all datasets. For the continuous treadmill perturbations [27], we divided the stance duration into 16 bins and computed a linear model for the ankle moment at each of those phases, all with delayed COM kinematics as inputs.

For each dataset, we pooled the data for the different perturbation magnitudes, perturbation direction (forward and backward), repetitions (i.e. trials) to perform a single least squares regression. The uncentered coefficient of determination (*R*^2^) (equations 7) and root mean square error (RMSE) between the measured and reconstructed joint moments and muscle activity are reported to quantify the fit of the linear regression (equations 1 and 2).

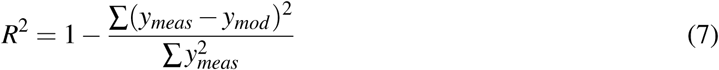

with *y*_*meas*_ the measured ankle moments or muscle activity and *y*_*mod*_ the reconstructed ankle moments and muscle activity with delayed linear feedback of COM kinematics.

### 4.5 Gains normalized by COP position or vertical ground reaction force

The hypothesis that gain modulation might be driven by cutaneous information from the foot was evaluated by fitting one model with constant feedback gains through all data points of the continuous perturbations (i.e. combing the different gait phases). We computed the *R*^2^ and *RMSE* when the feedback gains were proportional to the vertical ground reaction force (equation 5) or by the minimal distance between the COP and the bounds of the foot (equation 6) in the fore-aft direction. The bounds of the foot were computed based on the motion capture data.

### 4.6 Statistical analysis

We used statistical tests to evaluate (1) if gain modulation might be driven by cutaneous information from the foot and (2) if the fit with the task-level feedback model is better than the joint-level feedback model. To test the first hypothesis, we used a repeated measures ANOVA to evaluate if the *R*^2^ value is significantly different in models with constant gains modulated by the vertical ground reaction force or COP position compared to a model with constant gains (table 3). Bonferroni correction was used to correct for multiple comparisons in the post-hoc testing. To test the second hypothesis, we also used a paired t-test to evaluate if the *R*^2^ value is significantly different in the model with task-level feedback compared to a model with joint-level feedback. A two-sided confidence interval with an aplha level of 0.05 was used for all statistical tests.

## 5 Appendix

### 5.1 Time-delay sensitivity

We evaluated whether the variance in ankle moment explained by the linear regression is sensitive to the time delay. This additional analysis was important to verify that the observed relation between COM kinematics and ankle moment is a feedback control process, and does not simply reflect the coupling due to the dynamics of the skeleton. We did this sensitivity analysis on the pelvis push perturbations and found that the relation between ankle moment and COM kinematics is indeed sensitive to the neural delay (Fig. 11). The variance explained by the linear regression was optimal with a physiological plausible delay of 100ms and decreased strongly for delays smaller than 50 ms or larger than 120 ms. Hence, the observed correlation does not simply reflects the skeleton dynamics.

### 5.2 Perturbation magnitude: deviation in COM kinematics

To evaluate if the magnitude of the perturbations are similar between datasets, we compared the deviation in COM position and velocity at 150ms after perturbation onset for the four datasets (Fig. 12). We found that the deviations in COM position and velocity are similar in the perturbations during walking and are larger in perturbed walking compared to perturbed standing.

### 5.3 Statistical comparison feedback models

We evaluated if the feedback models with constant feedback gains modulated by (1) the vertical ground reaction force or (2) the position of the COP increased the *R*^2^ values compared to a model with only feedback gains. We used a paired t-test on the data of walking with continuous perturbations at three different speeds (Matlab 2019). We found only 1 significant result: For walking at 1.2 m/s the modulation of constant feedback gains with the vertical ground reaction force resulted in an significant increase in the *R*^2^ values compared to the model with constant gains (table 3).

## 6 Acknowledgments

The authors would like to thank the authors of the public available datasets [27] [9] for sharing their data. In addition, we would like to thank Herman van der Kooij and Edwin van Asseldonk for their insightful discussion related to the modulation of COM feedback during the gait cycle and if this modulation can be explained by altered input from tactile sensors.

